# *In situ* structure of the AcrAB-TolC efflux pump at subnanometer resolution

**DOI:** 10.1101/2020.06.10.144618

**Authors:** Muyuan Chen, Xiaodong Shi, Zhili Yu, Guizhen Fan, Irina Serysheva, Matthew L. Baker, Ben F. Luisi, Steven J. Ludtke, Zhao Wang

## Abstract

In Gram-negative bacteria, tripartite efflux pump AcrAB-TolC plays a prominent role in antibiotic resistance. We have used high resolution cryo-ET to visualize the structure of *Escherichia coli* AcrAB-TolC at a 7 Å resolution in intact cells. The resulting structures show the detailed architecture of the assembled complex embedded into cell envelope. Interactions with the inner membrane enable crosstalk between AcrB and TolC through AcrA, suggesting that assembly in the native cellular environment is critical for the pump activation mechanism, where the allosteric activating signal is triggered by the alternate binding of AcrA to the lipid membrane and AcrB porter domain. We establish a platform for high resolution *in situ* structural studies of bacteria efflux pump, which can yield critical information in understanding complex assemblies’ function.

**One sentence summary:** A 7 Å *in situ* structure of the AcrAB-TolC pump from bacteria by cryo-ET reveals mechanisms for active efflux.

---

Antibiotic resistance is an emerging crisis in healthcare as bacteria develop resistance to many widely used antibiotics. Bacteria efflux pumps, which actively expel a broad range of toxic substances, play a major role in the intrinsic and acquired drug resistance (*1*). In Gram-negative bacteria, efflux pumps are multicomponent assemblies that span the inner and outer membranes (*2*). In this study, we focus on *Escherichia coli (E. coli)* tripartite efflux pump AcrAB-TolC, which is comprised of the inner membrane transporter AcrB, the outer membrane channel TolC, and the periplasmic AcrA.

Previously, we have determined *in vitro* structures of AcrAB-TolC complex using cryo-EM single particle analysis (SPA) (*3*). These maps produced atomic models of the entire pump assembly, providing important mechanistic insights into the pump. However, since the AcrAB-TolC assembly spans the entire cell envelope, *in vitro* structures are far from native conditions. Thus, high-resolution structures within the cell would greatly aid in understanding how the native bacterial environment impacts the components and how they allosterically communicate.

Our previous *in situ* structures of the pump appear to be closed in the presence of antibiotics, and open when bound to inhibitors (*4*). The purified TolC is in a closed state (*5*), but TolC is open in the purified AcrAB-TolC complex in the presence of antibiotics or inhibitors (*3, 6*). In the absence of ligands, the TolC-closed complex can only be achieved using disulfide-engineered AcrA-AcrB crosslinked pump (Table S1) (*3*). The pumps’ different behavior *in vitro* and *in situ* suggests that the interactions between the pump and its surrounding membrane environment may play a critical role in activating the efflux process.

We have now resolved a 7 Å resolution structure of the AcrAB-TolC complex within *E. coli* cells in the presence of MBX3132 (an AcrB inhibitor) by cellular electron cryotomography (cryo-ET). Through a combination of improved sample preparation, data collection parameters, new subtomogram averaging software (*7*), and collection of a much larger data set, it was possible to achieve subnanometer resolution and provide a detailed view of the functioning pump in its native environment. This structure has sufficient resolution to observe the interactions of the pump with both membranes in much more detail and reveals sufficient secondary structural elements for accurate docking of high-resolution models. This new structure shows how interactions with the native membrane break the 6-fold symmetry, and lead to allosteric channel opening.

From the reconstructed tomograms, densely arranged AcrAB-TolC pumps can be clearly identified, where top view particles appear as small circles and side view particles appear as long thin channels spanning the cell envelope (Fig. 1). From 102 tomograms, we manually selected 27,932 particles located on the envelope of intact cells. Following *de novo* initial model generation, the structure of the fully assembled pump was determined to 7 Å resolution with C3 symmetry using the subtilt refinement pipeline in EMAN2 (*7*) (Fig. S1). Our previous *in situ* structure showed a large proportion of assembly intermediates (*4*), however subtomogram classification analysis of this new data does not show a clear sub-population of intermediate assembly states, confirming the function of MBX3132 in promoting and stabilizing pump assembly. The pump forms an open conduit from the funnel domain of AcrB toward the tip of TolC (Fig. 2A, C), which validates our previous prediction that MBX3132 will lock the assembled pumps in transporting state *in vivo* due to the saturation of binding pockets (*4*). At this resolution many individual alpha helices of AcrA and TolC are clearly resolved (Fig. 2A, B), permitting accurate docking of high-resolution structures. The bacteria peptidoglycan layer shows a weak density consistent with the location suggested in our previous *in situ* structures (*4*). Compared to the SPA structure of the AcrAB-TolC treated with inhibitor (PDB: 5NG5) (*3*), the helices in TolC and the upper part of AcrA match well with the *in vitro* pump structure, but the membrane proximal (MP) domain of AcrA in the cryo-ET structure is rotated 8 degrees (Fig. 2B). This rotation is likely caused by the AcrA termini being in its native environment, anchored into the inner membrane, rather than bound to free surfactant in the *in vitro* system. This represents a critical fact for understanding the function of the pump.

**Fig. 1.**
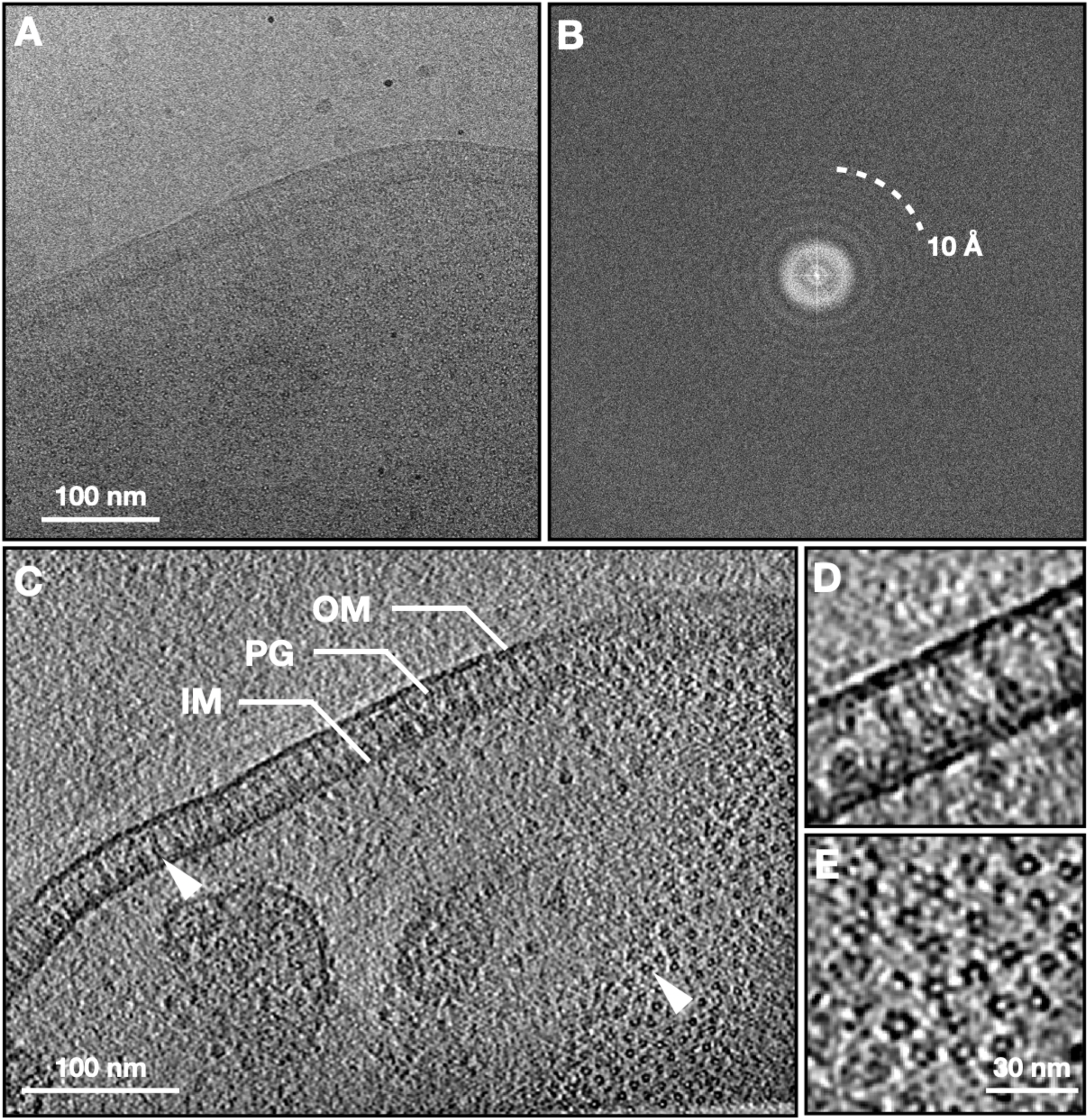
Cryo-ET of *E. coli* expressing the AcrAB-TolC multidrug efflux pump. (A) Zero-degree tilt image of a representative tilt series. (B) Fourier transform of (A). (C) Slice view of the reconstructed tomogram, with arrowheads showing the side and top view particles. (D, E) Zoomed in view of the side and top view particles. OM, outer membrane; IM, inner membrane; PG, peptidoglycan.

**Fig. 2.**
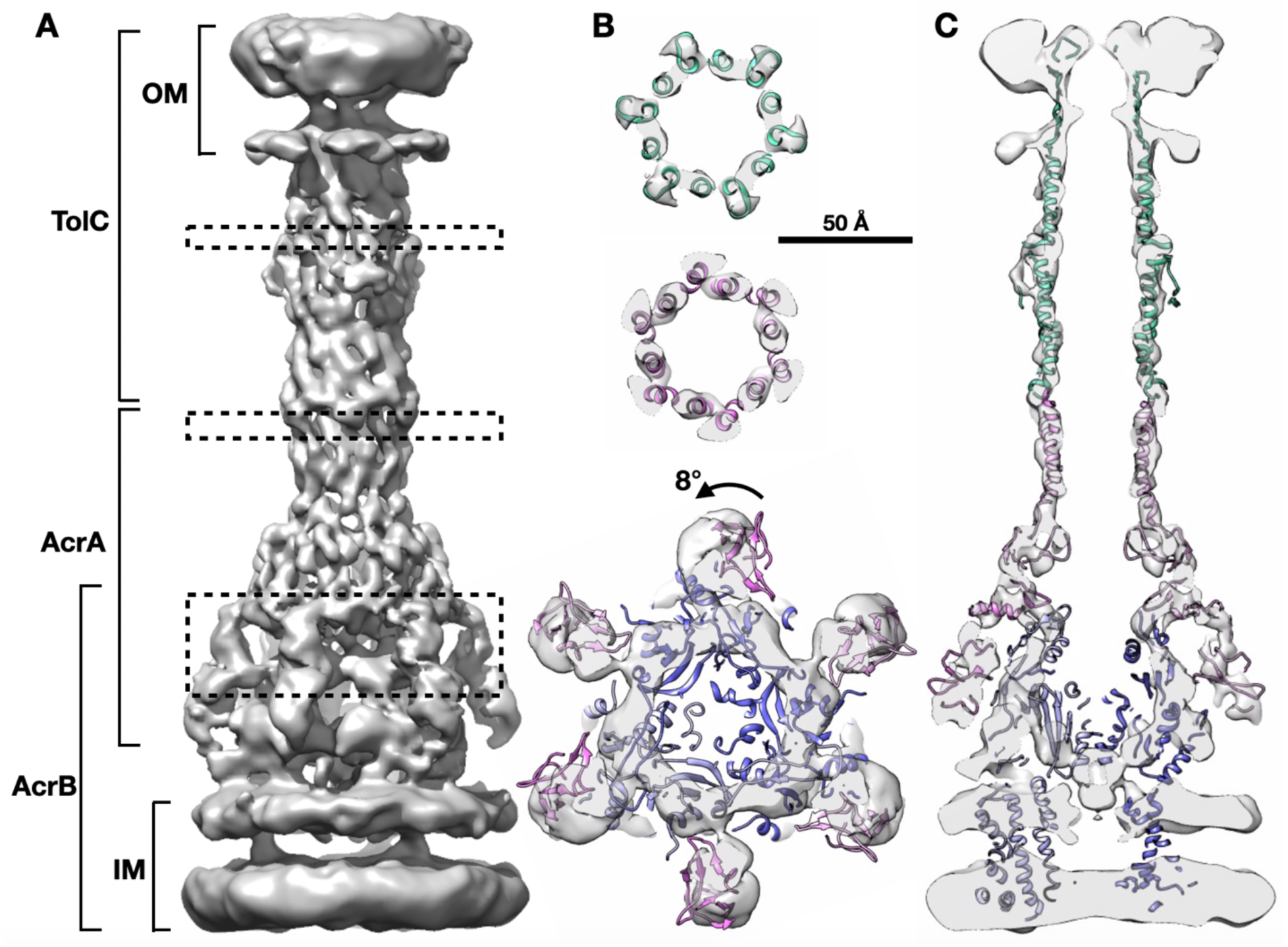
*In situ* structure of the AcrAB-TolC multidrug efflux pump. (A) Side view of the efflux pump density map. (B) Cross section views of the pump at the dashed box positions in (A), with the model of *in vitro* pump (PDB: 5NG5) fitted in density. (C) Cross section view through the center of the pump with the fitted model. OM, outer membrane; IM, inner membrane.

The density map is best resolved near the AcrA-TolC interface, but the resolution is worse in AcrB and the upper portion of TolC (Fig. S1). Although the structure of the complex was determined with C3 symmetry, TolC equatorial domains together with AcrA hexamer show a near C6 symmetry, suggesting that the map may be a mixture of conformations. To test this hypothesis, we performed focused refinements of the TolC region and the AcrB region respectively, with symmetry relaxed from C6 to C3. This resulted in separate averaged maps of the upper and lower parts of the pump.

In the focused refinement of TolC, the outer membrane beta-barrel, and equatorial helices are better resolved (Fig. 3A). In the density map, the outer leaflet of the outer membrane is much thicker than the inner leaflet, indicating the presence of the polysaccharide layer above the outer membrane. The loop at the top of the TolC beta-barrel is buried inside the polysaccharide layer (Fig. 3A), suggesting a critical role of the loop in expelling the polysaccharide and keeping an open path across the outer membrane. It has been suggested by several studies that the interiors of outer membrane proteins are often partially or completely occluded (*8*–*11*). As a result, previous crystal structures suggested that these loops have conformational mobility based on the sequence and may close over as a partial external ‘lid’ (*5*). According to our structure, the outer mouth is open with a diameter >10 Å, enough to expel substrates (Fig. 3A).

**Fig. 3.**
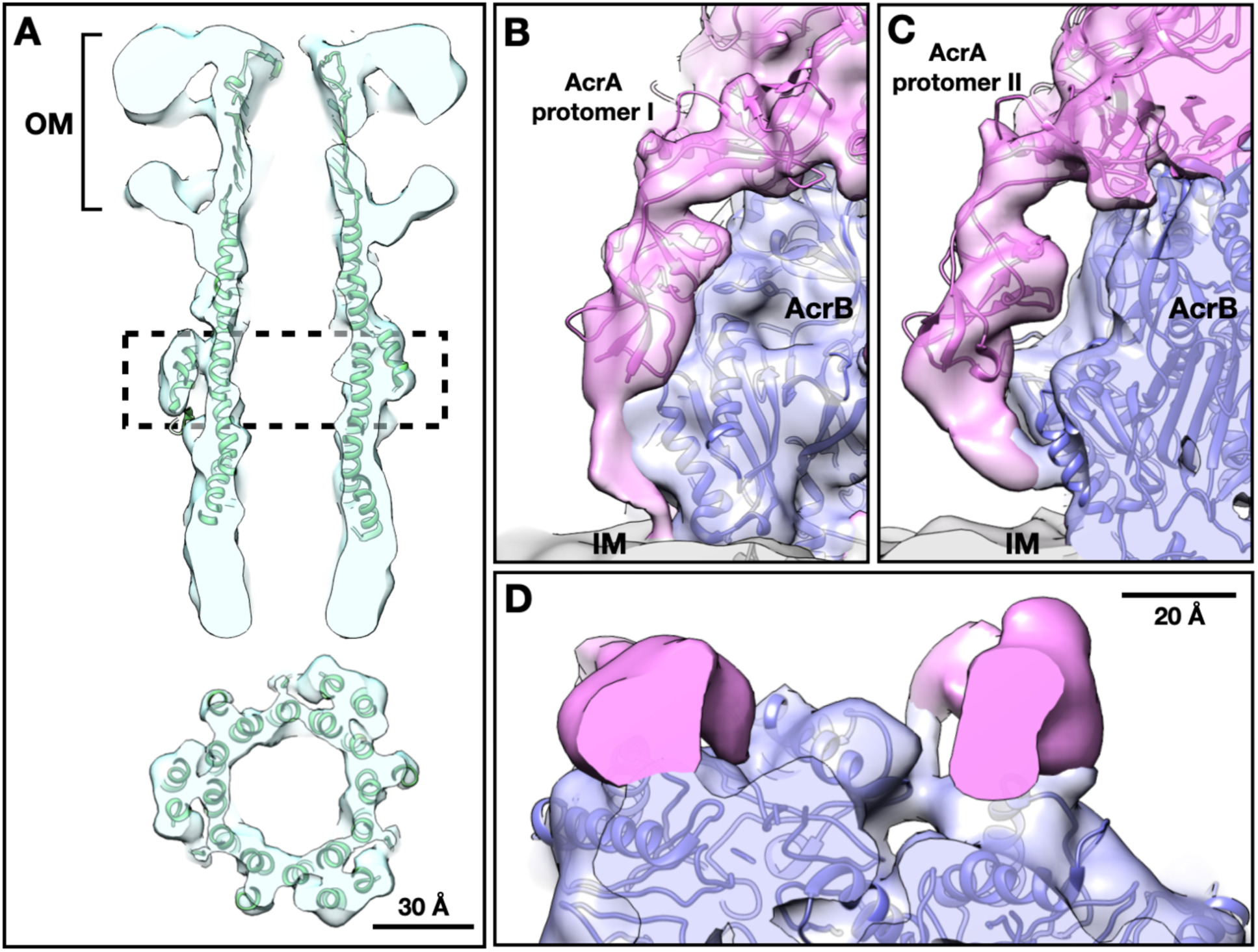
Focused refinement results. (A) Cross section views of TolC focused refinement result, with the model of *in vitro* pump (PDB: 5NG5) fitted in density. (B, C) Side view of the two protomers of AcrA terminus (pink) from the focused refinement result of the lower part of the pump. (D) Top view of the terminus of AcrA protomer I on the left and II on the right. OM, outer membrane; IM, inner membrane.

The TolC, AcrA, and AcrB form complex in a 3:6:3 ratio (*3*), which immediately leads to two possible relative orientations of TolC to AcrB, with a relative 60-degree rotation. Both possible complexes are observed in our data in a ∼1:1 ratio (Fig. S2), indicating that AcrAB subcomplex does not have a preference in recruiting the TolC in either orientation. These two different conformations of the overall complex also likely result in subtly different interactions between all three elements. This agrees with earlier findings for the homologous MexAB-OprM by SPA (*12*).

Compared to the *in vitro* structures of the entire pump, the focused refinement of the lower part of the pump shows a clear C3 symmetry. Densities of the transmembrane helix bundles are visible in the map. Notably the density of the AcrA terminus, which was not visible in previous structures, appears and strongly breaks the 6-fold symmetry (Fig. 3B-D). The density representing the six AcrA termini can be seen forming two distinct conformations interacting differently with AcrB and the membrane (Fig. 3B-D). The existence of a clear AcrA membrane anchor in only alternating subunits has not been previously hypothesized or observed, and in fact, could only be observed using the current *in situ* strategy. *De novo* models for the AcrA termini fit well into these densities, and hydrophobic residues near the N-terminus are expected to interact with the membrane (Fig. S3). These Rosetta models do not consider the surrounding membrane and protein densities, but still provide a conceptual description of our interpretation of the structure. In these models, the AcrA protomer I terminus closely interacts with the groove of the porter domain of AcrB (G639-S656) and is anchored into the inner membrane. The AcrA protomer II terminus sits between two AcrB subunits, forming solid contact with one AcrB subunit at the loop near S319. This terminus also interacts with the neighboring AcrB’s porter domain (PC2 subregion) (S834-K850).

AcrA interacts with AcrB and anchors in the inner membrane on one end, while connecting TolC and the peptidoglycan layer on the other end. This architecture allows AcrA to bridge and transfer conformational changes in AcrB in the inner membrane to TolC in the outer membrane. Compared to the *in vitro* models, AcrA in the *in situ* structure exhibits more extensive interactions with AcrB, and the newly revealed interaction sites are close to the substrate binding pockets of AcrB (Fig. 4). The AcrB porter domain, which interacts the AcrA terminus through PC2 subregion *in situ*, has the largest movement when binding substrates (*3*). Our model suggests that the interaction between the AcrA protomer II termini and the AcrB PC2 region in the porter domain, as well as the inner membrane anchoring of AcrA protomer I may work together to initialize the conformational change transition after the ligand binding at AcrB (Fig. 4).

**Fig. 4.**
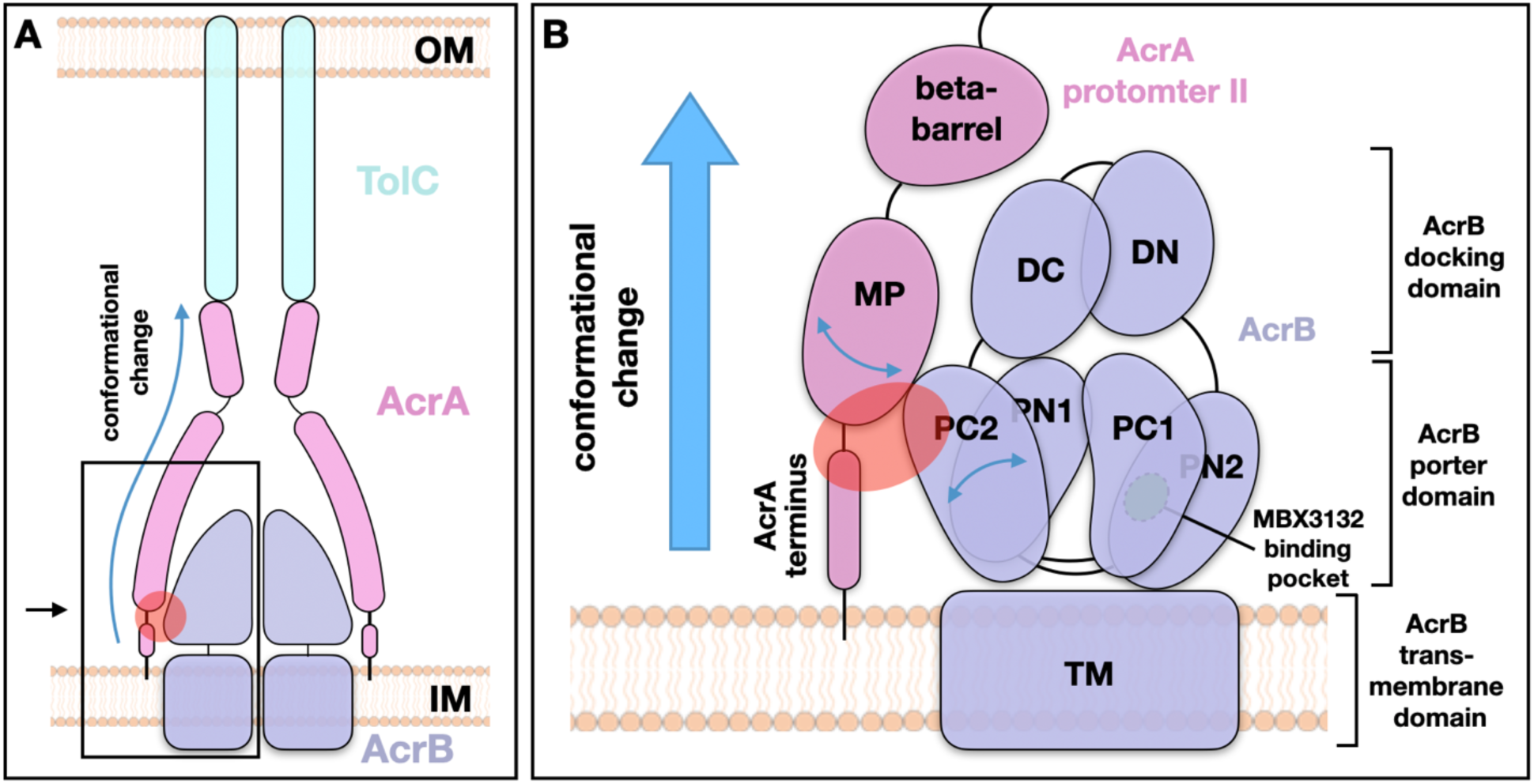
Interactions between AcrA and AcrB enable transmission of conformational changes from AcrB to TolC during substrate transportation. (A) Schematic view of the AcrAB-TolC efflux pump. (B) Zoomed in view of the boxed area in (A), viewing in the black arrow indicated direction. The red oval indicates the interaction zone between AcrA protomer II and AcrB. Substrate efflux from AcrB results in greatest movements in the PC2 domain. The MP domain of AcrA protomer II, which is anchored into the inner membrane, connects to the AcrB PC2 domain and can be affected by PC2 movement. The ensuing conformational change can be communicated through AcrA to the bottom of TolC to affect the gating of the channel. OM, outer membrane; IM, inner membrane.

Isolated TolC assumes a closed state (*3, 5*) before being recruited by AcrA to prevent leakage from the cell to the outside. Opening of TolC is required to form a continuous conduit through the periplasmic space to allow antibiotics efflux. We propose a model wherein substrate binding in AcrB triggers a conformational change in the porter domain which is propagated to TolC through the observed connections with AcrA. This movement in the porter domain will drive AcrA to transmit a twist toward the upper region and start to open the TolC channel through the hairpin interactions (Fig. 4).

In summary, our subnanometer *in situ* structures clearly demonstrates the importance of cellular environment in maintaining the proper structural interface and demonstrates the ability of cellular cryo-ET to retrieve critical structural data for understanding efflux activity. *In situ* structures fill in a critical knowledge gap of the tripartite pump assembly, and provide new insights into the mechanism of antibiotic resistance. The new AcrA-AcrB interaction sites revealed by the *in situ* structures which provide new insights into this major working mechanism of efflux pump, may also lead to a new direction in the future for drug development.

## Acknowledgements

We thank X. Yu, T. Huo and H. Wu for suggestions on sample preparation; T. Dendooven and A. Kirykowicz for helpful comments; and S. Raveendran for data backup. We thank Tim Opperman and colleagues for the kind gift of the AcrB inhibitor.

## Author contributions

Z.W. designed the experiments; S.J.L. developed computational methods. X.S. prepared and screened the sample; X.S., Z.Y. and G.F. performed data collection; M.C. performed computational analyses; M.L.B. performed model building. X.S., M.C., Z.Y., M.B. and Z.W. wrote the manuscript; G.F., I.S., B.F.L., S.J.L. and Z.W. reviewed and edited the manuscript.

## Data and materials availability

Density maps are available at EMDB with accession codes xxx. Atomic models are available in Protein Data Bank with accession codes xxx. Other data supporting this study is available from the corresponding author upon reasonable request.

## Supplementary Materials for

### Materials and Methods

#### Plasmid construction and protein expression

Plasmids pAcBH and pRSF-*tolC*, which carry the *acrAB* and *tolC* genes respectively, are preserved in our lab (*4*). To overexpress AcrAB-TolC multidrug efflux pumps, *E. coli* BL21 (DE3) cells (Invitrogen) were co-transformed with pAcBH and pRSF-*tolC*. Cells were cultured in 2xYT medium with antibiotics (100 *μ*g/ml ampicillin and 50 *μ*g/ml kanamycin) at 37 °C until an OD_600_ of 0.8 was reached and then induced by addition of 0.1 mM isopropyl β- d-1- thiogalactopyranoside at 20 °C overnight.

#### Sample preparation

*E. coli* cultures were harvested and washed by PBS buffer, then resuspended to an OD_600_ of 1.0. Cells were mixed with MBX3132 (1.4 *µ*g/ml) and incubated at 37 °C for 0.5 h. Subsequently, cells were mixed with 6 nm BSA fiducial gold (Aurion) and then deposited onto freshly glow-discharged, continuous floating carbon film covered grids (Quantifoil Au R3.5/1, 200 mesh). The grids were blotted with filter paper and plunge frozen in liquid ethane by using a Vitrobot Mark IV (FEI). Grids were stored in liquid nitrogen until data collection.

#### Cryo-ET data collection

Similar to our previous experiments (*4*), the cryo-ET data acquisition uses a bidirectional tilt scheme. The frozen-hydrated samples were imaged on a 300 kV Titan Krios transmission electron microscope (FEI) using a K2 Summit direct electron detector camera (Gatan), with a magnification of 81,000x. The pixel size is calibrated to be 1.67 Å. The tilt-series images were acquired at -1 to -2 *µ*m defocus range with an average cumulative dose of ∼90 e^−^/Å ^2^ distributed over 34 images and covering an angular range of -51 to +51 degrees, with an angular increment of 3 degree. 192 tilt series were collected in a 4-day imaging session.

#### Cryo-ET Data processing

First, movie frames of each individual tilt image are aligned using MotionCorr2. The tilt series alignment and tomogram reconstruction are performed using the automated workflow in EMAN2. Using the same workflow, defocus is determined for each tilt image, and CTF correction is performed at a per-particle-per-tilt level during particle extraction.

For the refinement of the full pump, 27,932 particles of the efflux pump are selected manually from 102 tomograms. The particles are then extracted from the tilt series with a box size of 256. 15 iterations of subtomogram refinement, followed by 4 iterations of subtilt refinement are performed using those particles and an initial model directly generated from the particles. The refinement process follows the “gold-standard” procedure, where particles are split into two independent subsets and aligned using different reference models that are phase randomized to 20 Å. At each iteration of the subtomogram refinement step, the worst 30% of the particles are excluded based on their similarity to the averaged structure. In the subtilt refinement step, sub-tilt images beyond 45 degrees are excluded, so are the worst 20% of the sub-tilt images. During the subtomogram refinement, the large-scale geometry information of the cells is used to guide the alignment and prevent the particles from being aligned up-side-down. Because the pump particles are tightly packed on the cell envelope, we simply draw a vector from the center of all particles in a tomogram to each particle, and rotate a particle by 180 degrees around the x axis if the particle is facing the opposite direction of the vector.

After subtomogram refinement of the full pump, particles of the upper and lower parts of the pump are re-extracted from the tilt series with a box size of 128, using the alignment information of the full pump. 10 iterations of subtomogram refinement are performed for each type of particle, using the corresponding part from the full pump structure as the initial model. The orientation search during the refinement is limited to 30 degrees from the Euler angle assigned in the full pump refinement. A symmetry breaking from C6 to C3 is also performed during the refinement. That is, for each particle after each round of the refinement, the similarity scores at the current orientation and at the C2 symmetrical orientation are compared, and the particle is assigned to its best orientation.

Since the entire refinement follows the “gold-standard” procedure (*13*), the reported resolution is measured by the spatial frequency that the Fourier Shell Correlation (FSC) curve between the masked structures determined from the two subsets of particles dropping below .143. Local resolution is measured by tiling the 3D structure with small overlapping cubes and calculating the FSC using the same criteria.

#### Model building

Modeling is first performed by rigid body fitting the models from high resolution cryo-EM structures into the cryo-ET density map using the Fit in Map tool in UCSF Chimera (*14*). To visualize the domain rotation of the full pump, the fitting focuses on the helical hairpin domain of AcrA. To show the interface between AcrA terminus and AcrB, the model of AcrA membrane proximal domain and AcrB are fitted into the focused refinement result respectively, and densities corresponding to the AcrB model are masked out from the averaged structure.

To further analyze the conformational differences between the *in situ* and *in vitro* pump structures, we flexibly fit our previous single particle structure of AcrAB-TolC complex with MBX3132 (PDB: 5NG5) into the *in situ* density map. First, the multimeric AcrA, AcrB and TolC portions of the 5NG5 were extracted and fit directly to the *in situ* map using UCSF Chimera. Once fitted, the individual complexes were then serially flexibly fit to the density map using the simulated annealing option in Phenix’s real space refinement tool, as well as FlexEM (*15*). Fittings with Phenix and FlexEM gave similar results. The individual complexes were then combined back into a single model and iteratively refined to maximize both fit to density and stereochemistry using real space refinement from Phenix (*16*) and manual optimization with Coot (*17*). As a note, the beta-sheet domain of TolC was not refined to the density due to the lack of features in this region. Rather, the final model contains the TolC beta-sheet extracted from 5NG5 concatenated to the refined portion of TolC. The final model was then assessed using MolProbity (*18*) and fit to density.

**Fig. S1.**
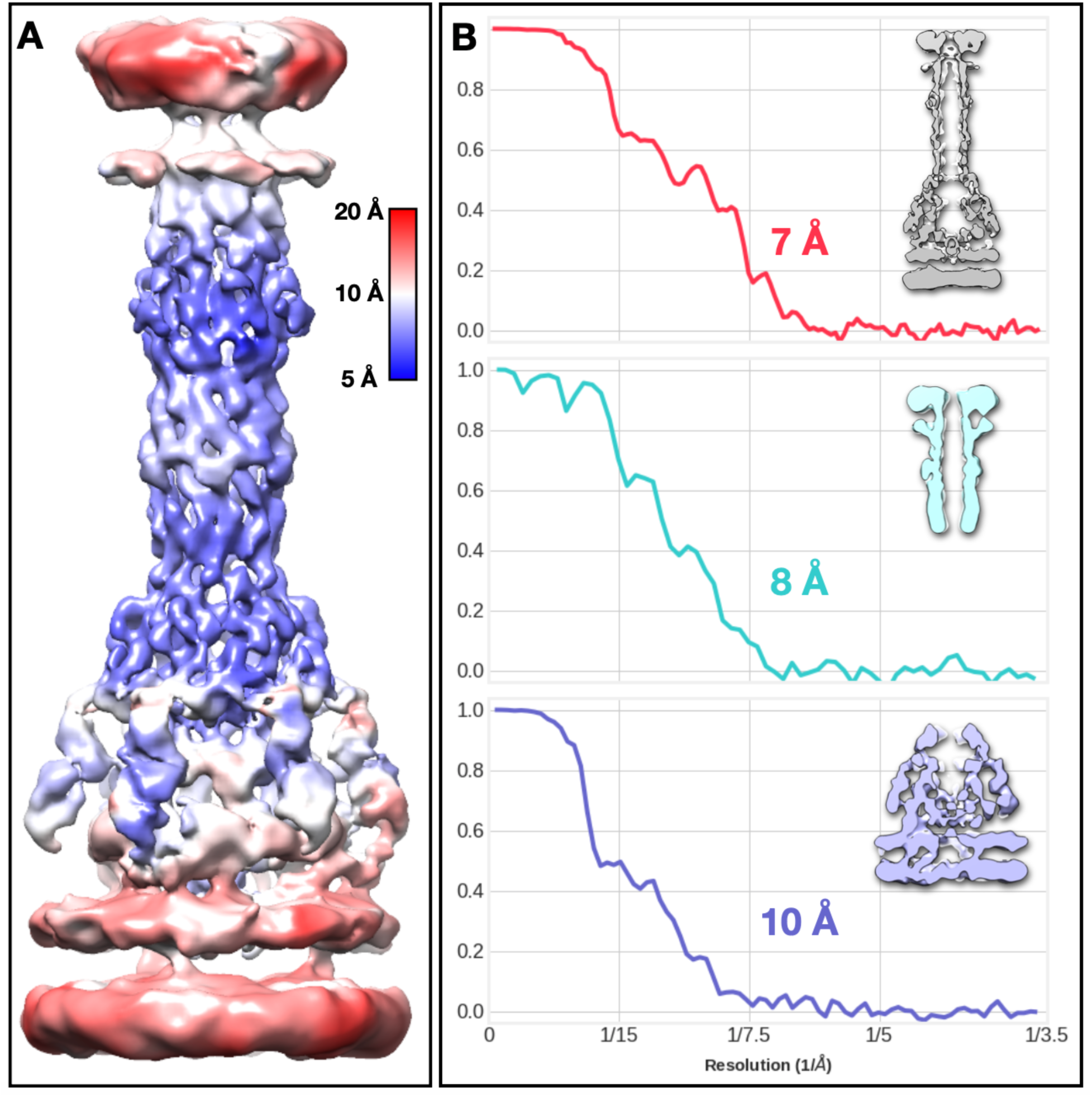
Local resolution estimate of subtomogram averaging results, using windowed FSC. (A) Averaged structure of the full pump colored by local resolution. (B) Fourier shell correlation plot of the full pump structure (7 Å, red); and the focused refinement results: TolC (8 Å, cyan); AcrB (10 Å, purple).

**Fig. S2.**
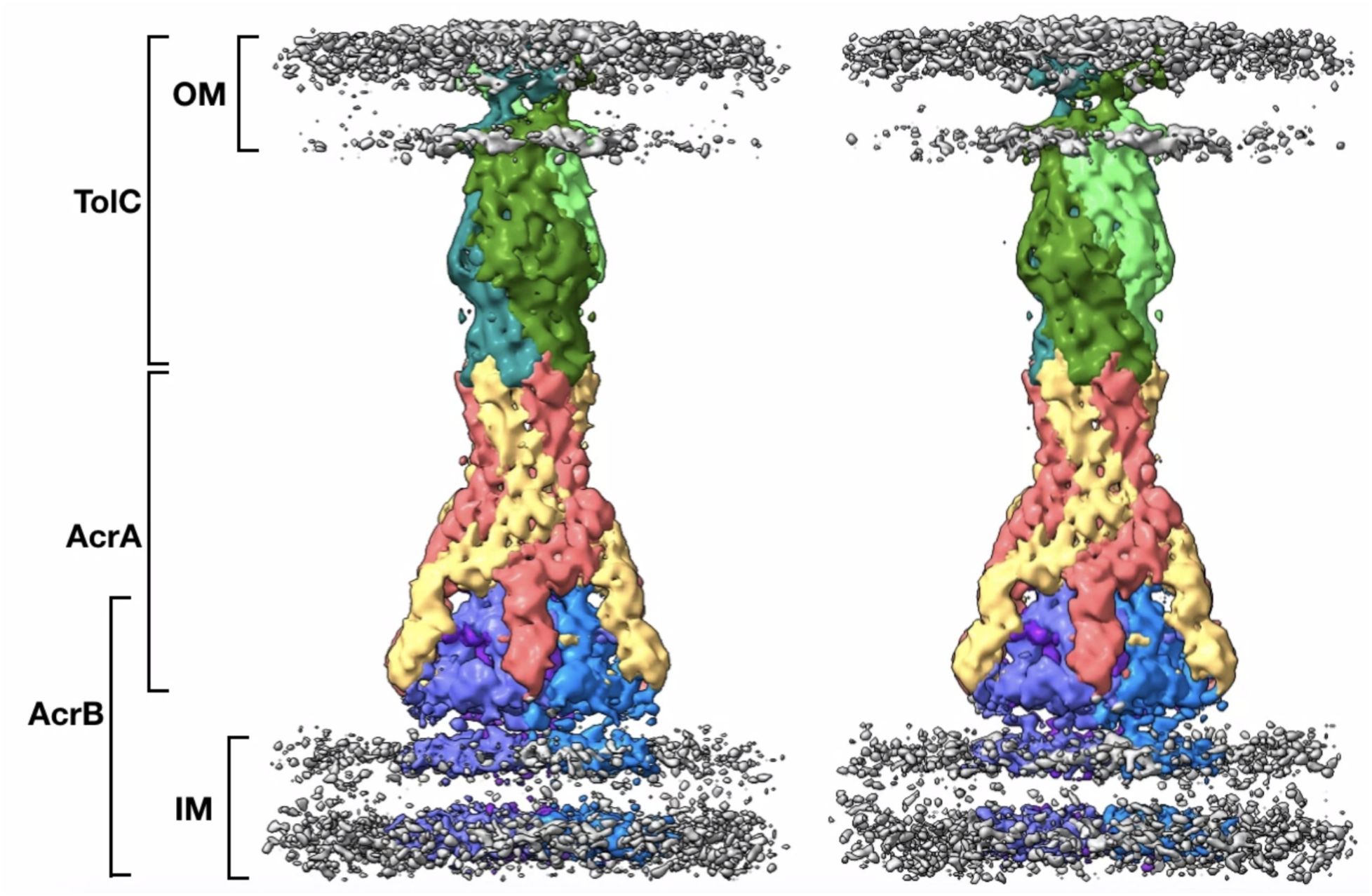
The two density maps generated separately based on the classification of TolC, representing two assembly models of the AcrAB-TolC efflux pump. Each monomer of TolC is notated by different but similar colors, so as each monomer of AcrB. AcrA protomer I and protomer II are notated by yellow and red colors, respectively. TolC has a 60-degree rotation relative to the AcrAB subcomplex. OM, outer membrane; IM, inner membrane.

**Fig. S3.**
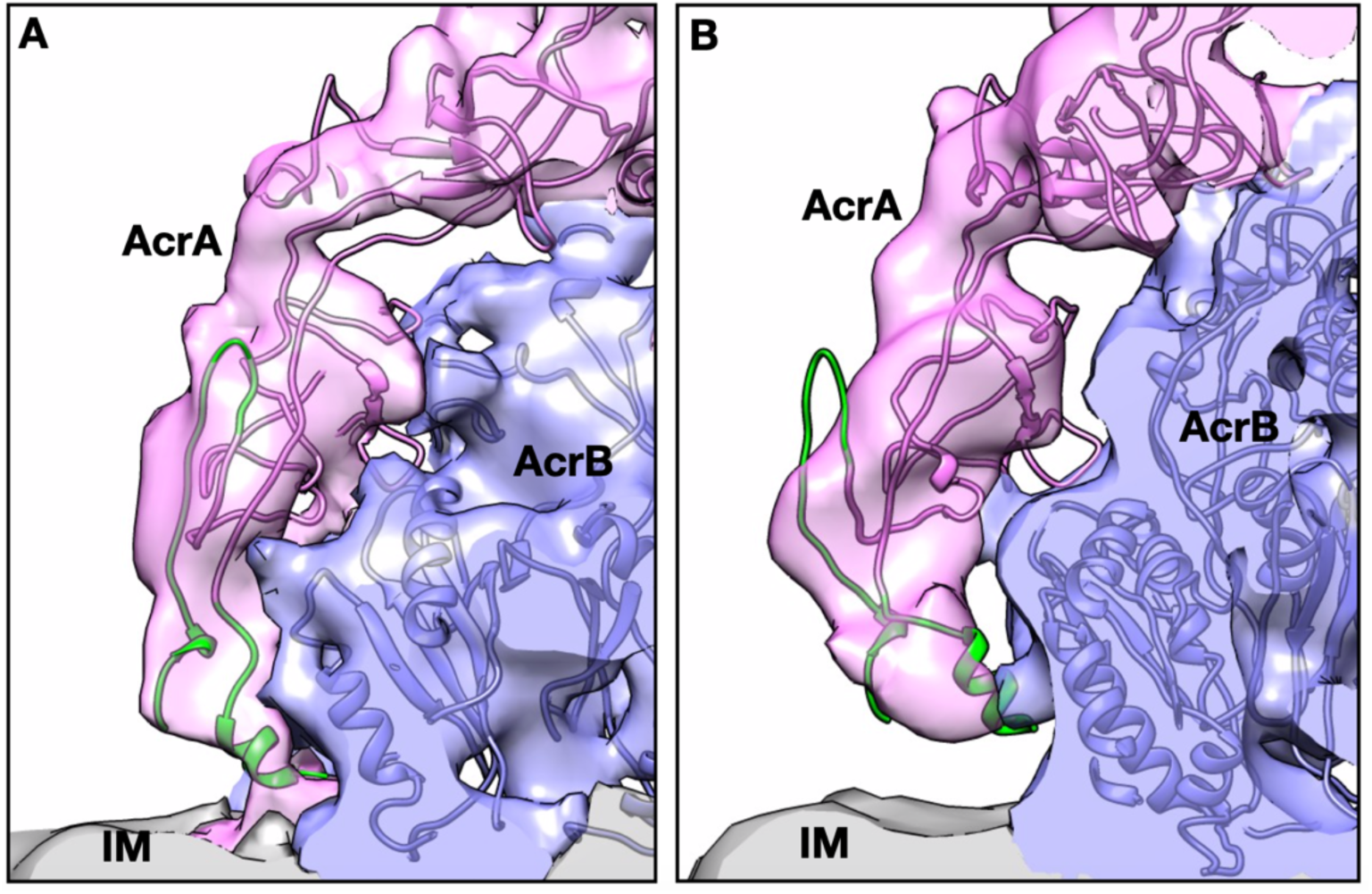
Side view of the interactions between AcrA and AcrB. (A) shows AcrA protomer I; and (B) shows AcrA protomer II. The *de novo* model for the AcrA termini colored green. IM, inner membrane.

**Table S1.** List of TolC channel states in different structural models from previous studies or this study. Inhibitor: MBX3132 (binds to AcrB and lock the pump at transporting state); antibiotics: puromycin; crosslinked AcrAB-TolC: the full pump assembly stabilized through the disulfide-bond linked AcrAB, generated by introducing cysteine-substitutions (AcrA-S273C and AcrB-S258C).

| Environment | Pump state | TolC channel state |
| --- | --- | --- |
| *in vitro* | isolated TolC (PDB: 1EK9) | closed |
|  | full pump+inhibitor (PDB: 5NG5) | open |
|  | full pump+antibiotics (PDB: 5O66) | open |
|  | crosslinked AcrAB-TolC (PDB: 5VS5) | closed |
|  | full pump apo state (EMD-5915) | open |
| *in vivo* | isolated TolC | N/A |
|  | full pump+inhibitor (this study) | open |
|  | full pump+antibiotics (EMD-0532) | closed |

**Movie S1. Visualization of the AcrAB-TolC efflux pump *in situ*, from cellular tomograms to averaged structures**.

